# New insight on DNA isolation from single automated recovered *Ceratocystis platani* conidia

**DOI:** 10.1101/2024.05.20.594938

**Authors:** Nicola Luchi, Pamela Pinzani, Francesca Salvianti, Irene Mancini, Alberto Santini

## Abstract

Single-cell technology is increasingly used to analyze the basis of molecular regulation and provide insights into different aspects of human diseases. Such technology is a breakthrough approach to study blood cancers by characterizing molecular information on a genome-wide scale at the single-cell level. These methods can be easily and successfully transferred to tracheomycotic plant pathogens, which cause host wilt. Ceratocystis platani is the causal agent of the Canker stain disease of plane tree (Platanus spp.), a lethal wilt disease spreading in Europe. To displace and separate different C. platani conidia types a dielectrophoretic approach was tested. The DNA of each conidium was isolated and analyzed and the target DNA was identified by a specific qPCR marker and by sequencing the amplicon. Our results showed that this technology is applicable to vascular plant pathogens. The fungal DNA was successfully extracted from single or pooled conidia and identified by both methods after whole genome amplification. The use of the single-cell technology will provide a new approach to the study of plant vascular diseases, allowing the study of single-spore molecular and physiological features not detectable in complex biological mixtures.

**Author summary:** In recent years, technologies for single-cell isolation have been developed in the study of human diseases, such as cancers, capable of obtaining genetic information at the single-cell level. In this study, these methods were transferred to a plant pathogen, Ceratocystis platani, which causes a lethal disease of plane tree. The single cell technique used was able to separate the different types of conidia of C. platani and analyze the DNA within each conidium. The use of single cell technology represents an important tool for the study of plant vascular diseases by allowing the study of molecular mechanisms that are difficult to detect in complex biological matrices.

## Introduction

The comprehensive analysis of DNA in complex biological mixtures has remained challenging. In recent years, rapid developments and technological improvements in high-throughput sequencing have enabled several applications with significant cost reductions. This, together with methodological advances, especially in the field of single-cell isolation, has paved the way for very reliable high-throughput analyses based on single cells.

Single-cell analysis allows for discovery of mechanisms undisclosed by the study of the whole population. It can be studied through the use of different technologies that allow metabolomic, proteomic, genomic and transcriptomic analysis at the single cell level [1–3]. Several techniques were developed for the separation of single cells from biological mixtures, including laser micro-dissection, fluorescence-activated cell sorting (FACS), and manual micromanipulation [4–6].

In clinical disciplines, such as cancer research, stem cell biology, immunology, developmental biology, and neurology single-cell analysis is very important to answer previously unsolvable questions; for example, the application of single-cell analysis in cancer research is expanding rapidly and promises to shed light on how a single cell can become metastatic or lead to cancer progression [7–10]. In recent years, technical advances have enabled molecular analyses at the single-cell level, allowing the profiling of rare cancer cells in clinical samples [11–12]. Single-cell analysis has been mainly devoted to the isolation of single cells from heterogeneous mixtures deriving from both cell lines and primary tissues; however, recently many efforts have been dedicated to the study of rare cell populations such as ‘Circulating Tumor Cells’ (CTCs) in the blood. In this case, despite the complexity of the experimental approach, the feasibility of single cell analysis has been demonstrated by several authors (e.g. [13–16]) relying on different workflows, and it is likely to have an impact on three major fields of oncology: early cancer detection and diagnosis, evaluation of disease progression, and prediction of therapeutic efficacy.

Most molecular techniques have been transferred from the clinical field to the field of plant pathology. The sensitivity of these approaches allows pathogens to be detected even before symptoms had developed on the plant [17–18]. The use of these new approaches based on single-cell analysis may also be relevant to study cell biology in plant-pathogenic microorganism such as fungi and bacteria. In previous generations, techniques for isolating unicellular forms of fungi and bacteria have been developed. The purposes of these isolations have been to study the life history, polymorphism, physiology, pathogenicity and finally the taxonomical classification of micro-organisms [19]. Spore and conidium are respectively units of sexual or asexual reproduction necessary for the dispersal and survival (often for long periods of time) of a microorganism. Often these propagules are carried by wind, insects, or may be present in other more or less complex matrices, such as soil, woody tissues, and water.

The causal agents of tracheomycosis, develop their mycelium and their propagules within the plant’s vascular tissue, causing vascular wilt disease on hundreds of plant species [20]. These pathogens have adapted to thrive in the xylem, which is a nutrient-poor niche. Recognition of vascular wilt pathogens by both extra-and intracellular plant receptors triggers plant innate immune responses that comprise physical and chemical defences [21]. Both types of defence responses occur in the xylem vessels in a coordinated manner, where physical defence responses mainly prevent the pathogens from spreading in the xylem vessels and chemical defence responses kill the pathogen or inhibit its growth. Sometimes the defences put in place by the plants leads to vessel obstructions drastically reducing hydraulic conductivity in the functional xylem, resulting in a severe wilt syndrome [22].

*Ceratocystis platani* is the causal agent of canker stain, a lethal disease of plane trees. This ascomycete fungus is native to North America and was introduced to Europe during World War II [23–25]. The pathogen is now present in France, Switzerland, Italy, Albania, Greece [26] and European Turkey [27] in both urban and natural environments. Oriental and London plane trees are at risk of extinction in both urban and natural areas, as there are no limiting factors to the spread of this pathogen wherever plane trees grow [28].

The fungus enters the host through wounds and then moves through the xylem as a wilt pathogen, but also kills cambium and colonizes parenchymatic rays [26].*Ceratocystis platani* is mainly transmitted by human activities such as pruning and terracing, and is naturally transmitted via root anastomosis, through infected water and via the frass of *Platypus cylindrus*, a beetle commonly present on dying plane trees [29].

*Ceratocystis platani* isolates are self-fertile and produce, often within 10 days, many fruiting bodies (ascomata) on the surface of infected wood or in culture medium. Ascomata are dark brown to black and globose with a long thin neck, through which the ascospores are exuded. They accumulate in a sticky drop at the top of the ascomatal stalks, where they appear as a cream-coloured mass [26]. Three forms of conidia are produced by the asexual stage of the fungus: cylindrical and doliform endoconidia and globose to pyriform aleuroconidia (**Fig 1**). It is believed that the smell produced by these droplets, and the attractive sugary substances within them, represent adaptations favouring dispersal by insects [30].Pure isolation of individual spores or conidia represents a challenge in the study of microorganism biology, especially in the host tissues. An advantage of studying single units is that the presence of possible inhibitors in the matrices in which propagules are included would be significantly reduced. The possibility of investigating these reproductive and spreading structures through transcriptomics would clarify either the existence of a differential role between the sexual and asexual structures, or between different forms of conidia. Last but not least, it would be possible to study their movement within the host tissues and especially vessels.

**Fig 1.**
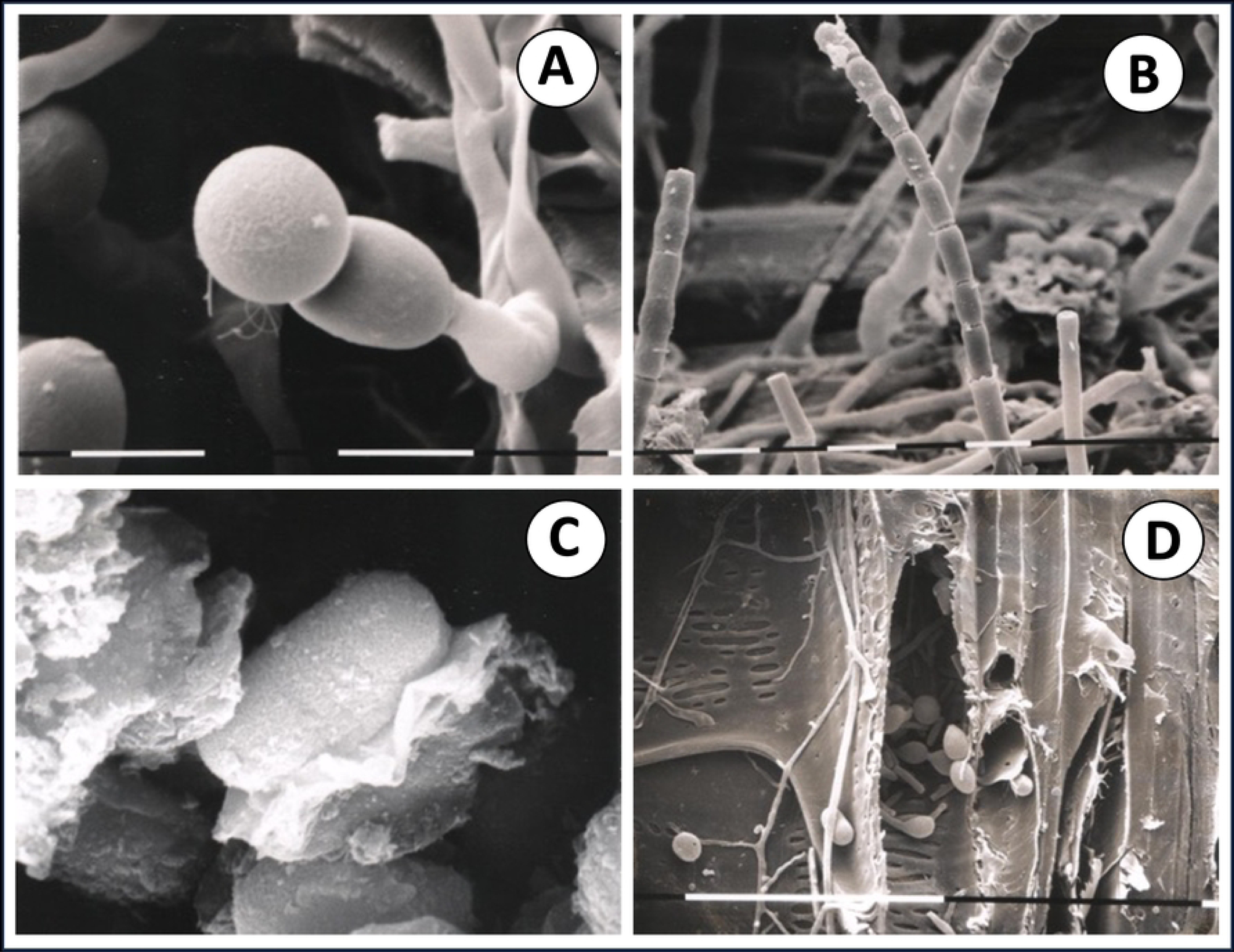
Scanning Electron Microscope (SEM) images of *Ceratocystis platani* propagules and reproductive structures. A) aleuroconidia (x 2,500); B) cylindrical endoconidia (x1,250); C) ascospore (x10,000); D) germinating conidia within host vascular tissue (x505; 100μm) (Photo by Dr. Alberto Panconesi, CNR - Firenze).

Among the various methods of single-cell analysis reported, dielectrophoresis (DEP) has been the one recently acquired for the isolation of single cells. Dielectrophoresis refers to the motion of a polarizable particle when subjected to a nonuniform electric field due to the interaction between the spatial gradient of the electric field and the dipole of the particle. Particle movement is influenced by the ambient electric field and by the properties of dielectric particle or solutions [31–33]. Because biological cells have different dielectric properties, dielectrophoresis can be used to sort, transport and separate different cell types; thus achieving a single-cell level of purification. Dielectrophoresis has been applied in several industrial fields, i.e., microfluidics, biosensors, medical diagnostics, and environmental studies [34–37], as well as in medical science to study cancer cells, virus, bacteria and fungi [38,15,16].

Application of the DEP technique has not been applied in plant pathology [39]. To date the only use of DEP was those described by [40] that developed a portable microfluidic device employed for the dielectrophoresis-driven capture of spores and subsequent on-chip detection of *Sclerotinia sclerotiorum* airborne inoculum. The use of DEP in the plant pathology field may shed light on unknown mechanisms of the biology of plant pathogens. For these reasons, the aim of this work was to set up and validate an active automated trapping tool to recover *C. platani* single conidia and to isolate DNA by using this new approach based on dielectrophoresis systems (**Fig 2**).

**Fig 2.**
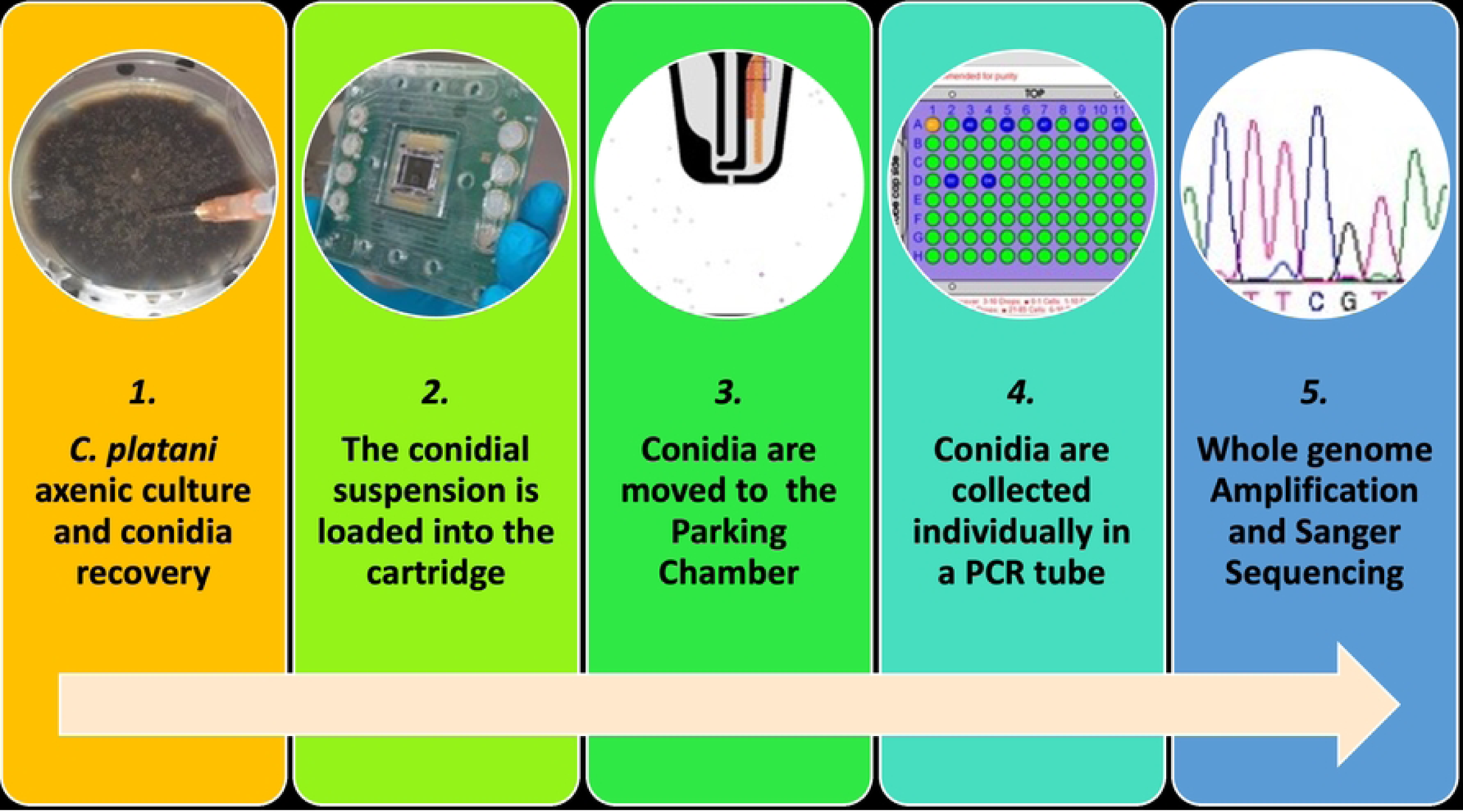
Schematic representation of the entire workflow for the molecular analysis of a single conidium.

## Results

### Fungal conidia recovery

*Ceratocystis platani* conidia developed on 15-day-old axenic culture were stained with DAPI and recovered and sorted according to their different shapes using DEPArray system. Each conidia selected population (aleuroconidia, cylindrical endoconidia, and clustered aleuroconidia) were collected and analyzed (**Fig 3**). The number of the whole *C. platani* population conidia, dispensed in the main chamber, was 10,616. Among them, a total of 290 differently shaped conidia were isolated: i) 90 single cylindrical endoconidia (oblong shaped), and ii) 180 single aleuroconidia (globular shaped). In addition, 20 samples were also obtained from clustered aleuroconidia (**Fig 4a**). Each cluster contains from 5 to 15 conidia and the size is 60μm or less. This size has been evaluated considering that each square of the DEPArray cage is approximately 60 x 60μm (**Fig 3**). Using DEPArray™, images of individual conidia showed positive fluorescent signals for DAPI, allowing easy detection and recovery of different conidia shapes.

**Fig 3.**
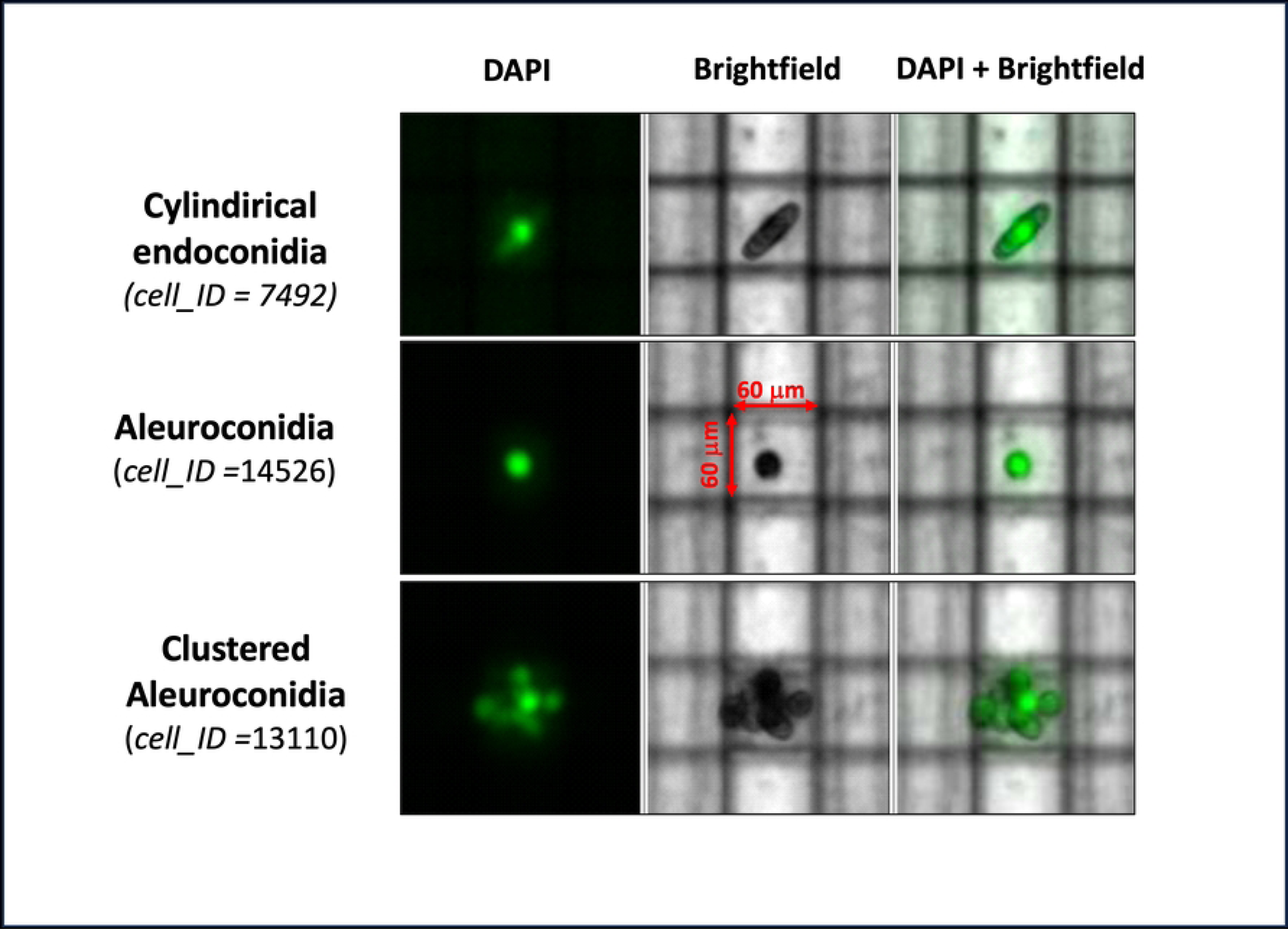
Different conidia shapes of *Ceratocystis platani* as imaged by the DEPArray system (DAPI positive fluorescent signal and brightfield). Images of: i) cylindrical endoconidium (ID 7492), ii) aleuroconidium (ID 14526) and iii) clustered aleuroconidia (ID 13110) are shown in the figure.

**Fig 4.**
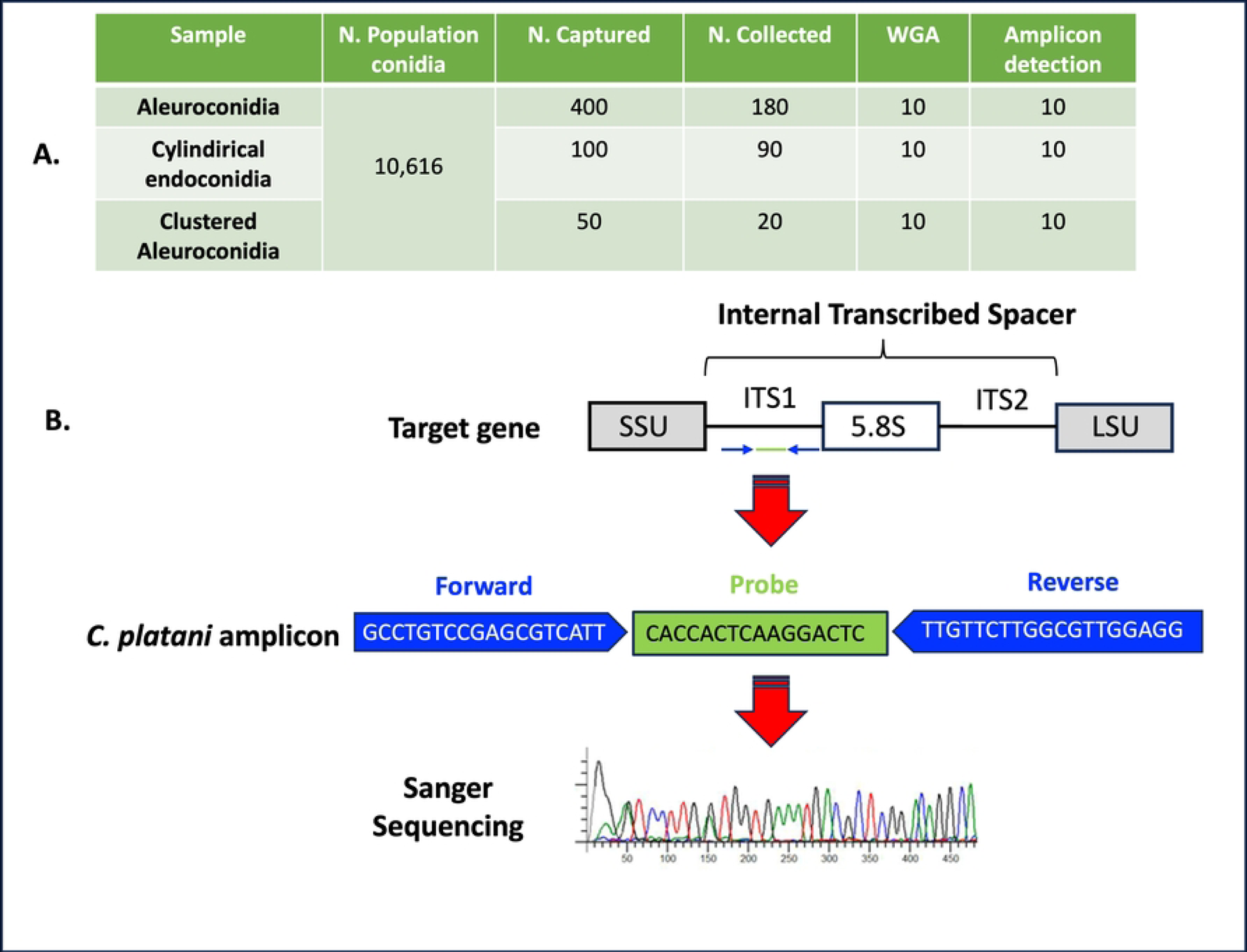
*Ceratocystis platani* conidia collected and analyzed by using DEPArray™. A) Different shaped conidia isolated. The number of population conidia is referred to conidia present in the main chamber; B) qPCR ITS Amplicon target [41] sequenced from single conidia.

### DNA amplification from conidia

The purity of *C. platani* conidia recoveries has been demonstrated by molecular analysis. As a proof of concept, we sequenced Ampli1 WGA products obtained from the 10 randomly chosen individual conidia (**Fig 4a**), and in all cases the system was able to read the sequence of the ITS amplicon (as previously reported by [41] (**Fig 4b**). The specificity of the *C. platani* amplicon was also confirmed by the nucleotide–nucleotide BLAST ® search showing a complete homology (100%) between the *C. platani* amplicon sequences designed in [41] and the *C. platani* sequences available in the NCBI database. These results proved that the single cell method was able to isolate individual *C. platani* conidia and amplify their DNA.

## Discussion and conclusions

Through the DEPArray™ microchip, a state-of-the-art automated technology, it was possible for the first time to recover and analyze DNA from a single conidia of *C. platani*, a fungal pathogen of plane trees. Usually, genomic analysis of single cells is performed using sequencing technologies such as whole genome sequencing (WGS) and RNA sequencing [42,43]. Single-cell sequencing allows measurement of cell-to-cell variations that are normally masked by conventional sequencing, where each sequencing library is represented by a population of cells instead of a single cell [43]. Single-cell sequencing has been applied in different fields of clinical research, for example to study the copy number variation between tumoral and healthy cells [44,45], recombination activity in human sperm [46] and to identify gene expression dynamics among different cell subpopulations [47]. More recently, single-cell genomic techniques represent a powerful approach to study tumors at single-cell resolution, providing a novel approach to deciphering cancer biology [48]. This technology has been enormously successful for the isolation of CTCs in blood [6,16], as by exploiting electric fields, it allows individual cells to be moved directly into microtubes so as to analyze their cell genome in purity. Although the field of single-cell plant genomics is at its initial stages, there is great potential for obtaining information on cell lineage and functional cell types to understand more accurately complex cellular interactions in plant tissue [49].

By using the DEPArray technology, the operator can select and isolate cells (conidia) with a well-defined shape and size. This is important for fungal microorganisms because they do not have only one type of cell morphology, but can have different forms that allow them to acquire new virulence properties in the host, becoming successful pathogens [50]. In the mammalian host, changes in morphology represent a common and effective strategy for pathogen survival; thus avoiding the attack of the immune system. Indeed, some studies have shown that during interactions with the host, human pathogenic fungi go through a series of morphological changes that are closely associated with virulence [51,52]. However, these transitions are also important in plant pathogenic fungi to cause disease. Studies have shown that different fungal species with genetically identical spores, formed at different developmental stages or environmental conditions, exhibit different physical and physiological properties [53,54]. Kang et al. [55], performed a single-cell study on *Aspergillus fumigatus* and found that germination phenotypes vary among identical individuals showing that the environment of spore production determines the size, the degree of spore germination and the impact virulence in the host. In addition, *Magnaporthe oryzae*, the causal agent of rice blast, undergoes a series of morphotype changes during the infection process [56], while *Ustilago maydis*, the pathogenic agent of corn smut, grows as yeast on artificial media and infects plants, establishing a biotrophic interaction with the host, only as filamentous dikaryon [57].

Recently, attention has also been paid to understanding the role of different forms of conidia belonging to a species. Many fungal species, pathogenic to plant species, have distinct asexual conidia. However, the role of these propagules often remains uncertain, and few studies have been conducted so far. A recent work by Nordzieke et al. [58], demonstrated how two different types of conidia (“falcate” and “oval”) of *Colletotrichum graminicola,* the causal agent of anthracnose on maize, showed differences in germination, early colony formation and infection processes, having distinct roles in pathogenesis. Despite the presence of several forms of *C. platani* propagules, *i.e*., ascospores, endoconidia, aleuroconidia, their distinct roles have not yet been clarified. Therefore, the use of the single-cell technique will be essential to better understand their role and function in the fungal infection process.

This innovative technology, exploiting principles of microelectronics and microfluidics, makes it possible to identify and separate different cell types using high-resolution fluorescence and bright-field images. The graphical interface allows researchers to select the desired cells easily and with accuracy. Visual identification of conidia is also done with nuclear markers such as DAPI (4′,6-Diamidino-2-phenylindole), that emit blue fluorescence when bound to DNA. The use of an intercalating dye has several benefits and allows selection and collection of the desired conidia while ensuring quality control of the samples, since broken nuclei and those with dissolved membranes are easily visible. In addition, debris is easily distinguished from labeled conidia and the pool of conidia.

One of the classic methods for quantitatively determining the number of cells (conidia or spores) in a suspension involves the use of a standard microbiological instrument (Burker Haemacytometer Counting Chambers) that, through mathematical formulas, provides an estimate of the total number of cells per unit of volume. One of the disadvantages of this method, is that cells are identified only by size, and it is possible that dust or other particles may cause a count error. The use of the single-cells technique is able to overcome these problems because, by using new technologies, it has made it possible for the first time to select either a single conidium or a well-defined number of conidia that make up the analyzed pool.

Through the use of the DEPArray the operator is able to display on the monitor the selected spores and then, thanks to the electromagnetic field present inside the microchip, is able to move the spores (single or pooled) into the microtube and use them for DNA or RNA release and further molecular analysis (**Fig. S1**). Determination of the exact number of cells is also important for precisely defining the standards that are used in the preparation of the calibration curve in quantitative PCR. In this way, corresponding amounts of DNA can be associated with an exact number of spores, allowing for greater precision in quantitative DNA measurement.

Microbial biodiversity due to the simultaneous presence of various fungal and bacterial species in plant tissues may cause contamination in pure colony isolation experiments. If conidial suspensions were prepared from these colonies, the latter could contain the conidia of other microbial species, or different strains of the same species; thus compromising the purity of the sample. Because each colony or strain can affect the growth of another, the ability to separate and culture axenic colonies, consisting of only one type of pathogen, is essential in diagnostic settings. Pure cultures allow for additional molecular and genetic analysis for identification and characterization of infectious agents.

In this context, the use of single-cell analysis also avoids possible interference between different populations of conidia, allowing only the conidia under analysis to be selected and sampled. In addition, through the electromagnetic field generated in the microchip, the selection of conidia also avoids possible interference with PCR inhibitors, which may be found within the fruit from which the suspension is obtained. For example, polyphenols and polysaccharides, which represent PCR inhibitors [59–61], are very common compounds in fungi [62, 63].

Although the field of single-cell plant genomics is at the beginning, there is great potential for obtaining information on cell lineage and functional cell types to understand more accurately the complex cellular interactions in plant tissue [64]. Fungi produce conidia that often remain dormant until environmental conditions are suitable for germination. These propagules contain abundant stable messenger RNAs. However, the mechanisms behind the production, composition and functions of these transcripts are still poorly understood. Recently, Wang et al. [65], found how genetically identical conidia mature into phenotypically variable conidia. Indeed, the authors have shown that fungal conidia synthesize and store transcripts according to the environmental conditions present before dormancy, thus preparing for the future. For this reason, the use of cutting-edge techniques allows for a better understanding of biological processes at the single-conidia level.

Through the DEPArray™ system, the biological processes of the isolated cells, individually contained in tiny droplets of buffer, remain unaltered, and the DNA and RNA are not fragmented but intact. This allows the highest quality genetic analysis to be conducted on isolated (single or pooled) cells. Genomic and transcriptomic analysis of a single conidium (or spore) provides new solutions for studying fungal microorganisms both before their colonization of the host (dormant spores or conidia) and during their colonization of tissues; such as in the case of *C. platani* where conidia (and spores) are also produced within the host plant vascular system. By studying single spores in plant cells or tissues, it will be possible to better understand the movement, development and biological activity of the pathogenic microorganism in plant tissues and the response of plants to these biotic stresses. Although there may be challenges in single-cell preparation, DNA/RNA amplification, DNA sequencing, and data analysis, the rapid evolution of single-cell technologies will play an important role in the study of plant-pathogen interactions; thus, enabling a better understanding of some biological mechanisms that are not yet understood.

## Materials and Methods

### Fungal conidia suspension

A *Ceratocystis platani* axenic culture (Cp26 isolates) was obtained from a symptomatic plane (*Platanus acerifolia*) tree. Fungal isolation was carried out from woody samples collected from the boundary between necrotic and healthy tissues. Briefly, small tissue fragments were removed from the margin of the necrotic portion, and placed in 90 mm Petri dishes (Sarstedt, Verona, Italy) on 1.5% potato dextrose agar (PDA; Difco Laboratories, Detroit, MI) and incubated at 20°C in the dark. After 15 days, a 1ml sterile syringe needle was scraped onto the mycelial colony and rinsed in a 1.5ml microtube (Sarstedt, Italy) containing 100μl sterile distilled water. The conidia suspension was adjusted to 10^5^ conidia ml^−1^ using a Burker chamber.

### *Ceratocystis platani* conidia isolation by using DEPArray^TM^ process

A drop of NucBlue™ Live Cell ReadyProbes™ Reagent (DAPI) (Life Technologies Italia, Monza, Italy), was added to 1ml of conidia suspension and incubated in the dark for 10 min. After centrifugation (800 rcf for 5 min) the supernatant was removed. The conidia were then washed in 1ml with SB115 buffer (Menarini Silicon Biosystems, Bologna, Italy) and centrifuged (800 rcf for 5 min). The entire buffer was then removed, taking care not to remove the pellet, resuspended in 13μl SB115 buffer and loaded onto a DEPArray^TM^ cartridge (Menarini Silicon Biosystems).

The operating principle of the DEPArray^TM^ is dielectrophoresis (DEP), an electrokinetic principle based on the ability of a non-uniform electric field to polarize single cells, which are trapped in a stable levitation, avoiding any contact between cells and surfaces during the capture process (Di Trapani et al., 2018). The isolation of single or pooled *C. platani* conidia was carried out using the DEPArray technology that includes: the single cell-sorting instrument, a microfluidic disposable cartridge (DEPArray^TM^ A300K DS V2.0) and the CellBrowser^TM^ software (Menarini Silicon Biosystems). This technology uses a microsystem cartridge, a disposable device which integrates a microelectronic silicon chip, microfluidic chambers and valves.

The architecture of the DEPArray™ microchip consists of three different chambers: 1) Main chamber; 2) Parking chamber, and 3) Exit Chamber [66]. Briefly, the ascospores suspension (in SB115 buffer) is loaded in the Main chamber of the cartridge (S1 **Fig**). The chip scanning is carried out by an automated fluorescence microscope in the instrument (DEPArray cell separator), generating an image gallery in the DEPArray system. In accordance with their morphology, single or clustered fungal conidia were selected in each ‘DEP cage’, and automatically moved in the Parking chamber (S1 **Fig**), where multiple groups of different conidia can be parked simultaneously before recovery. Then, the target single or pooled conidia are eluted from the device into 200 µL microtubes (S1 **Fig**). All samples were washed in phosphate-buffered saline (PBS) and then preserved at −80°C with 1 µL PBS, for subsequent analysis.

### Whole genome amplification and sequencing

To obtain a sample suitable for sequencing analysis, Whole Genome Amplification (WGA) of the genome of the single conidia was performed using the Ampli1™ WGA kit (Menarini Silicon Biosystems). The procedure is based on a ligation-mediated PCR following a site-specific DNA digestion by *MseI* enzyme. The kit has no need for precipitation steps, avoiding DNA loss, and uses a mixture of Taq polymerase with a proofreading enzyme, *Pwo* polymerase, that has been reported to have error rates more than 10 times lower than the error rate observed with *Taq* polymerase [67]. As the Ampli1™ WGA kit was optimized for human genomes, we verified the impact of the site specific (AATT) DNA digestion on the genome of *C. platani* to highlight the possible impact of the procedure on the target of the subsequent sequencing analysis. We also checked the possible presence of these specific sites on a target *C. platani* gene (ITS -Internal Transcribed Spacer) where qPCR primers and probes were designed [41].

The complete amplification of the total DNA present in each single conidia (or conidia cluster) was performed using the Ampli1™ WGA Kit (Menarini Silicon Biosystems) following manufacturer instructions. Amplified DNA from ascospores was used for ITS target gene analysis which was performed by Sanger sequencing. Briefly, the target region in the ITS gene was amplified using *C. platani* specific forward and reverse primers CpITS-F (GCCTGTCCGAGCGTCATT) and CpITS-R (CCTCCAACGCCAAGAACAA) as already reported by Luchi et al. [41]. The presence of the product of the amplification reaction was initially verified by melting analysis and PCR products were then sequenced using the BigDye^TM^ Terminator v1.1 Cycle Sequencing Kit (Life Technologies). The sequence reaction was purified using the ZR DNA Sequencing Clean-Up Kit (Zymo Research, California, USA) and analyzed using an ABI PRISM 310 Genetic Analyzer (Applied Biosystems, USA). The specificity of the amplicon was checked in silico using the basic local search tool (BLAST) in the NCBI (National Center for Biotechnology Information) database.

## Acknowledgments

The Authors wish to thank Dr. Matthew Haworth, Institute for Sustainable Plant Protection (IPSP-CNR), for his help in language manuscript editing. This study was funded by European Commission Horizon 2020 Research and Innovation Programme ‘HOlistic Management of Emerging forest pests and Diseases’ (HOMED) (grant No 771271). This work was partly funded by the “Accordo di collaborazione per attività scientifica Regione Toscana - IPSP-CNR n. AOOGRT0365836-02/10/2019”.

## Authors contribution

**Conceptualization:** Nicola Luchi, Alberto Santini, Pamela Pinzani

**Data curation:** Nicola Luchi, Pamela Pinzani

**Formal analysis:** Pamela Pinzani, Francesca Salvianti, Nicola Luchi

**Funding Acquisition:** Alberto Santini

**Methodology:** Francesca Salvianti, Irene Mancini, Nicola Luchi, Pamela Pinzani

**Resources:** Pamela Pinzani, Alberto Santini

**Supervision:** Nicola Luchi, Pamela Pinzani

**Writing – original draft:** Nicola Luchi, Alberto Santini, Pamela Pinzani

**Writing – review & editing:** Nicola Luchi, Alberto Santini, Pamela Pinzani, Francesca Salvianti, Irene Mancini

## Supporting information captions

**S1 Fig. Single cell method based on the DEPArray™ microchip to isolate *C. platani* conidia.** A) Conidia (black dots) are loaded in the “Main chamber” (1), the electric field is able to move the conidia into the “Parking chamber” (A2) (B), where different clusters of conidia were grouped and each single conidium visualized with a different colored circle (red, orange or purple) (C). Each target conidium is then moved to the “Exit chamber” (A3), eluted in a single drop (arrow) of PBS buffer (D) and (E) automatically placed in a single microtube.

## Reference

1. Baysoy A, Bai Z, Satija R, Fan R. The technological landscape and applications of single-cell multi-omics. Nat Rev Mol Cell Biol. 2023; 24: 695–713. 10.1038/s41580-023-00615-w

2. Stuart T, Satija R. Integrative single-cell analysis. Nat Rev Genet. 2019; 20: 257–272. 10.1038/s41576-019-0093-7

3. Zhang K, Zemke NR, Armand EJ, Ren B. A fast, scalable and versatile tool for analysis of single-cell omics data. Nat Methods. 2024; 21(2):217–227. 10.1038/s41592-023-02139-9

4. Antoniadi I, Skalický V, Sun G, Ma W, Galbraith DW, Novák O, Ljung K. (2022). Fluorescence activated cell sorting-A selective tool for plant cell isolation and analysis. Cytometry. 2022; 101(9): 725–736. 10.1002/cyto.a.24461.

5. Fröhlich J, König H. New techniques for isolation of single prokaryotic cells. FEMS Microbiol Rev. 2000;24(5):567–72. 10.1111/j.1574-6976.2000.tb00558.x.

6. Pinzani P, D’Argenio V, Del Re M, Pellegrini C, Cucchiara F, Salvianti F, Galbiati S. (2021). Updates on liquid biopsy: current trends and future perspectives for clinical application in solid tumors. Clin Chem Lab Med. 2021; 59(7): 1181–1200. 10.1515/cclm-2020-1685

7. Choi JH, Lee BS, Jang JY, Lee YS, Kim HJ, Roh J, Shin YS, Woo HG, Kim CH. Single-cell transcriptome profiling of the stepwise progression of head and neck cancer. Nat Commun. 2023;14(1):1055. 10.1038/s41467-023-36691-x.

8. Grüntkemeier L, Khurana A, Bischoff FZ, Hoffmann O, Kimmig R, Moore M, Cotter P, Kasimir-Bauer S. Single HER2-positive tumor cells are detected in initially HER2-negative breast carcinomas using the DEPArray™-HER2-FISH workflow. Breast Cancer. 2022; 29(3):487–497. 10.1007/s12282-022-01330-8.

9. Lawson DA, Bhakta NR, Kessenbrock K, Prummel KD, Yu Y, Takai K, Zhou A, Eyob H, Balakrishnan S, Wang CY, Yaswen P, Goga A, Werb Z. Single-cell analysis reveals a stem-cell program in human metastatic breast cancer cells. Nature. 2015; 526 (7571):131–135. 10.1038/nature15260.

10. Lim B, Lin Y, Navin N. Advancing cancer research and medicine with single-cell genomics. Cancer cell. 2020; 37(4): 456–470. 10.1016/j.ccell.2020.03.008.

11. Hosseini H, Obradovic MMS, Hofmann M, Harper KL, Sosa MS, Werner-Klein M, Nanduri LK, Werno C, Ehrl C, Maneck M, Patwary N, Haunschild G, Guzvic M, Reimelt C, Grauvogl M, Eichner N, Weber F, Hartkopf AD, Taran F-A, Brucker SY, Fehm T, Rack B, Buchholz S, Spang R, Meister G, Aguirre-Ghiso JA, Klein CA. Early dissemination seeds metastasis in breast cancer. Nature. 2016; 540(7634): 552–558. 10.1038/nature20785.

12. Salvianti F, Orlando C, Massi D, De Giorgi V, Grazzini M, Pazzagli M, Pinzani P. Tumor-related methylated cell-free DNA and circulating tumor cells in melanoma. Front Mol Biosci. 2016; 2: 76. 10.3389/fmolb.2015.00076.

13. Lohr JG, Adalsteinsson VA, Cibulskis K, Choudhury AD, Rosenberg M, Cruz-Gordillo P, Francis JM, Zhang CZ, Shalek AK, Satija R, Trombetta JJ, Lu D, Tallapragada N, Tahirova N, Kim S, Blumenstiel B, Sougnez C, Lowe A, Wong B, Auclair D, Van Allen EM, Nakabayashi M, Lis RT, Lee GS, Li T, Chabot MS, Ly A, Taplin ME, Clancy TE, Loda M, Regev A, Meyerson M, Hahn WC, Kantoff PW, Golub TR, Getz G, Boehm JS, Love JC. Whole-exome sequencing of circulating tumor cells provides a window into metastatic prostate cancer. Nat Biotechnol. 2014;32(5):479–84. 10.1038/nbt.2892.

14. Rangel-Pozzo A, Liu S, Wajnberg G, Wang X, Ouellette RJ, Hicks GG, Drachenberg D, Mai S. Genomic Analysis of Localized High-Risk Prostate Cancer Circulating Tumor Cells at the Single-Cell Level. Cells. 2020;9(8):1863. 10.3390/cells9081863.

15. Pestrin M, Salvianti F, Galardi F, De Luca F, Turner N, Malorni L, Pazzagli M, Di Leo A, Pinzani P. Heterogeneity of PIK3CA mutational status at the single cell level in circulating tumor cells from metastatic breast cancer patients. Mol Oncol. 2015;9(4):749–57. 10.1016/j.molonc.2014.12.001.

16. Salvianti F, Pazzagli M, Pinzani P. Single circulating tumor cell sequencing as an advanced tool in cancer management. Expert Rev Mol Diagn. 2016; 16(1): 51–63. 10.1586/14737159.2016.1116942.

17. Luchi N, Capretti P, Pazzagli M, Pinzani P. Powerful qPCR assays for the early detection of latent invaders: interdisciplinary approaches in clinical cancer research and plant pathology. Appl Microbiol Biotechnol. 2016; 100: 5189–5204. 10.1007/s00253-016-7541-5

18. Luchi N, Ioos R, Santini A. Fast and reliable molecular methods to detect fungal pathogens in woody plants. Appl Microbiol Biotechnol 2020; 104 (6): 2453–2468. 10.1007/s00253-020-10395-4

19. Davis WH. Single Spore Isolation. Proc Iowa Acad Sci. 1930; 37(1):151–159.

20. Yadeta KA, Thomma BPHJ. The xylem as battleground for plant hosts and vascular wilt pathogens. Front Plant Sci. 2013; 4:97. 10.3389/fpls.2013.00097.

21. Agrios GN. Plant Pathology, 5th Edn. Amsterdam: Elsevier Academic Press, 2005

22. Santini A, Faccoli M. Dutch elm disease and elm bark beetles: a century of association. iForest. 2015; 8(2): 126–134. 10.3832/ifor1231-008

23. Panconesi A. Canker stain of plane trees: a serious danger to urban plantings in Europe. J Plant Pathol. 1999; 81(1): 3–15.

24. Cristinzio M, Marziano F, Verneau R. La moria del platano in Campania. Rivista di Patologia Vegetale.1973; 189–214.

25. Ferrari JP, Pichenot M. The canker stain disease of plane tree in Marseilles and in the south of France. Eur J For Pathol. 1976; 6(1): 18–25.

26. Tsopelas P, Santini A, Wingfield MJ, Wilhelm de Beer Z. Canker stain: a lethal disease destroying iconic plane trees. Plant Dis. 2017; 101(5): 645–658. 10.1094/PDIS-09-16-1235-FE

27. Lehtijärvi A, Oskay F, Doğmuş Lehtijärvi HT, Aday Kaya AG, Pecori F, Santini A, Woodward S. *Ceratocystis platani* is killing plane trees in Istanbul (Turkey). For Pathol. 2018; 48(1): e12375. 10.1111/efp.12375

28. Jeger M, Bragard C, Chatzivassiliou E, Dehnen-Schmutz K, Gilioli G, Jaques Miret JA, MacLeod A, Navajas Navarro M, Niere B, Parnell S, Potting R, Rafoss T, Urek G, Van Bruggen A, Van der Werf W, West J, Winter S, Santini A, Tsopelas P, Vloutoglou I, Pautasso M, Rossi V. Scientific opinion on the risk assessment and reduction options for *Ceratocystis platani* in the EU. EFSA Journal. 2016;14(12): 4640, 65 pp. 10.2903/j.efsa.2016.4640

29. Soulioti N, Tsopelas P, Woodward S. *Platypus cylindrus*, a vector of *Ceratocystis platani* in *Platanus orientalis* stands in Greece. For Pathol. 2015; 45(5): 367–372. 10.1111/efp.12176

30. Malloch D, Blackwell M. Dispersal biology of the ophiostomatoid fungi. In: Wingfield MJ, Seifert KA, Webber JF, editors. Ceratocystis and Ophiostoma. St. Paul, MN: APS Press; 1993. pp. 195–206.

31. Lee D, Hwang B, Kim B. The potential of a dielectrophoresis activated cell sorter (DACS) as a next generation cell sorter. Micro and Nano Syst Lett. 2016; 4:2. 10.1186/s40486-016-0028-4

32. Song H, Rosano JM, Wang Y, Garson CJ, Prabhakarpandian B, Pant K., Klarmann GJ, Perantoni A, Alvarez LM, Lai E. Continuous-flow sorting of stem cells and differentiation products based on dielectrophoresis. Lab Chip. 2015; 15(5): 1320–1328. 10.1039/c4lc01253d.

33. Alshareef M, Metrakos N, Juarez Perez E, Azer F, Yang F, Yang X, Wang G. Separation of tumor cells with dielectrophoresis-based microfluidic chip. Biomicrofluidics. 2013; 7(1): 011803. 10.1063/1.4774312.

34. Zhang J, Yan S, Alici G, Nguyen NT, Di Carlo D, Li W. Real-time control of inertial focusing in microfluidics using dielectrophoresis (DEP). Rsc Advances. 2014; 4(107): 62076–62085. 10.1039/C4RA13075H.

35. Tomkins MR, Chow J, Lai Y, Docoslis A. A coupled cantilever-microelectrode biosensor for enhanced pathogen detection. Sensors and Actuators B: Chemical. 2013;176: 248–252. 10.1016/j.snb.2012.09.020.

36. Švorc Ľ, Rievaj M, Bustin D. (2013). Green electrochemical sensor for environmental monitoring of pesticides: Determination of atrazine in river waters using a boron-doped diamond electrode. Sensors and Actuators B: Chemical. 2013; 181:294–300. 10.1016/j.snb.2013.02.036

37. Adekanmbi EO, Srivastava SK. Dielectrophoretic applications for disease diagnostics using lab-on-a-chip platforms. Lab on a Chip. 2016;16(12): 2148–2167. 10.1039/C6LC00355A.

38. Abd Rahman N, Ibrahim F, Yafouz B. Dielectrophoresis for Biomedical Sciences Applications: A Review. Sensors. 2017;17(3):449. 10.3390/s17030449.

39. Donoso A, Valenzuela S. In-field molecular diagnosis of plant pathogens: recent trends and future perspectives. Plant Pathol. 2018; 67(7):1451–1461. 10.1111/ppa.12859.

40. Duarte PA, Menze L, Shoute L, Zeng J, Savchenko O, Lyu J, Chen J. Highly efficient capture and quantification of the airborne fungal pathogen *Sclerotinia sclerotiorum* employing a nanoelectrode-activated microwell array. ACS Omega. 2021; 7(1): 459–468. 10.1021/acsomega.1c04878.

41. Luchi N, Ghelardini L, Belbahri L, Quartier M, Santini A. Rapid detection of *Ceratocystis platani* inoculum by quantitative real-time PCR assay. Appl Environ Microbiol. 2013;79 (17): 5394–5404. 10.1128/AEM.01484-13.

42. Kolodziejczyk AA, Kim JK, Svensson V, Marioni JC, Teichmann SA. The Technology and Biology of Single-Cell RNA Sequencing. Molecular Cell. 2015; 58(4); 610–620. 10.1016/j.molcel.2015.04.005.

43. Shapiro E, Biezuner T, Linnarsson S. Single-cell sequencing-based technologies will revolutionize whole-organism science. Nat Rev Genet. 2013;14(9):618–630. 10.1038/nrg3542.

44. Baslan T, Kendall J, Rodgers L, Cox H, Riggs M, Stepansky A, et al. Genome-wide copy number analysis of single cells. Nat. Protoc. 2012; 7:1024–1041.10.1038/nprot.2012.039.

45. Navin N, Kendall J, Troge J, Andrews P, Rodgers L, McIndoo J, et al. Tumor evolution inferred by single-cell sequencing. Nature. 2011; 472:90–94. 10.1038/nature09807.

46. Wang J, Fan HC, Behr B, Quake SR. Genome-wide single-cell analysis of recombination activity and de novo mutation rates in human sperm. Cell. 2012; 150:402–412. 10.1016/j.cell.2012.06.030.

47. Saliba AE, Westermann AJ, Gorski SA, Vogel J. Single-cell RNA-Seq: Advances and future challenges. Nucleic Acids Res. 2014; 42:8845–8860. 10.1093/nar/gku555.

48. Suvà ML, Tirosh I. Single-Cell RNA Sequencing in Cancer: Lessons Learned and Emerging Challenges. Mol Cell. 2019; 75(1):7–12. 10.1016/j.molcel.2019.05.003.

49. Yuan Y, Lee H, Hu H, Scheben A, Edwards D. Single-Cell Genomic Analysis in Plants. Genes 2018. 9(1):50. 10.3390/genes9010050.

50. Min K, Neiman AM, Konopka JB. Fungal pathogens: shape-shifting invaders. Trends in Microbiol. 2020; 28(11): 922–933. 10.1016/j.tim.2020.05.001.

51. Li Z, Nielsen K. Morphology changes in human fungal pathogens upon interaction with the host. J Fungi. 2017; 3(4):66. 10.3390/jof3040066.

52. Köhler JR, Hube B, Puccia R, Casadevall A, Perfect JR. Fungi that infect humans. Microbiol Spectr. 2017; 5(3). 10.1128/microbiolspec.funk-0014-2016.

53. Hagiwara D, Sakai K, Suzuki S, Umemura M, Nogawa T, Kato N, Osada H, Watanabe A, Kawamoto S, Gonoi T, Kamei K. Temperature during conidiation affects stress tolerance, pigmentation, and trypacidin accumulation in the conidia of the airborne pathogen *Aspergillus fumigatus*. PLoS One. 2017; 12(5):e0177050. 10.1371/journal.pone.

54. Oliveira M, Pereira C, Bessa C, Araujo R, Saraiva L. Chronological aging in conidia of pathogenic *Aspergillus*: comparison between species. J Microbiol Methods. 2015; 118: 57–63. 10.1016/j.mimet.2015.08.021.

55. Kang SE, Celia B, Bensasson D, Monamy M. Sporulation environment drives phenotypic variation in the pathogen *Aspergillus fumigatus*. G3 Genes Genom Genet. 2021;11(8). 10.1093/g3journal/jkab208.

56. Wilson RA, Talbot NJ. (2009). Under pressure: investigating the biology of plant infection by *Magnaporthe oryzae*. Nat Rev Microbiol. 2009;7(3):185–195. 10.1038/nrmicro2032.

57. Brefort T, Doehlemann G, Mendoza-Mendoza A, Reissmann S, Djamei A, Kahmann R. *Ustilago maydis* as a pathogen. Ann Rev Phytopathol. 2009; 47: 423–445. 10.1146/annurev-phyto-080508-081923.

58. Nordzieke DE, Sanken A, Antelo L, Raschke A, Deising HB, Pöggeler S. Specialized infection strategies of falcate and oval conidia of *Colletotrichum graminicola*. Fungal Genet Biol. 2019; 133:103276. 10.1016/j.fgb.2019.103276.

59. Schrader C, Schielke A, Ellerbroek L, Johne R. PCR inhibitors – occurrence, properties and removal. J Appl Microbiol. 2012; 113 (5): 1014–1026. 10.1111/j.1365-2672.2012.05384.x.

60. Jobes DV, Hurley DL, Thien LB. Plant DNA isolation: A method to efficiently remove polyphenolics, polysaccharides, and RNA. Taxon.1995; 44: 379–386. 10.2307/1223408.

61. Wilson IG. Inhibition and facilitation of nucleic acid amplification. Appl Environ Microbiol. 1997. 63(10): 3741–3751. 10.1128/aem.63.10.3741-3751.1997.

62. Jung JY, Lee IK, Seok SJ, Lee HJ, Kim YH, Yun BS. Antioxidant polyphenols from the mycelial culture of the medicinal fungi *Inonotus xeranticus* and *Phellinus linteus*. J Appl Microbiol. 2008; 104(6): 1824–1832. 10.1111/j.1365-2672.2008.03737.x.

63. Yang Y, Li J, Hong Q, Zhang X, Liu Z, Zhang T. Polysaccharides from *Hericium erinaceus* fruiting bodies: structural characterization, immunomodulatory activity and mechanism. Nutrients. 2022;14(18):3721. 10.3390/nu14183721.

64. Yuan Y, Lee H, Hu H, Scheben A, Edwards D. Single-cell genomic analysis in plants. Genes. 2018; 9(1): 50. 10.3390/genes9010050.

65. Wang F, Sethiya P, Hu X, Guo S, Chen Y, Li A, Tan K, Wong KH. (2021) Transcription in fungal conidia before dormancy produces phenotypically variable conidia that maximize survival in different environments. Nat Microbiol. 2021;6(8):1066–1081. 10.1038/s41564-021-00922-y.

66. Di Trapani M, Manaresi N, Medoro G. DEPArray™ system: An automatic image-based sorter for isolation of pure circulating tumor cells. Cytometry. 2018; 93: 1260–1266. 10.1002/cyto.a.23687.

67. McInerney P, Adams P, Hadi MZ. Error Rate Comparison during Polymerase Chain Reaction by DNA Polymerase. Mol Biol Int. 2014; 287430. 10.1155/2014/287430.

